# Reciprocal recurrent selection based on genetic complementation: An efficient way to build heterosis in diploids due to directional dominance

**DOI:** 10.1101/2022.07.05.498857

**Authors:** Giovanny Covarrubias-Pazaran, Christian Werner, Dorcus Gemenet

**Affiliations:** Excellence in Breeding Platform, Consultative Group of International Agricultural Research, Mexico; International Maize and Wheat Improvement Center (CIMMYT), Texcoco, Mexico

**Keywords:** Hybrid breeding, heterosis, molecular markers, alleles, haplotypes, quantitative trait locus

## Abstract

Depending on the trait architecture and reproduction system, selection strategies in plant breeding focus on the accumulation of additive, dominance effects, or both. Innovation in the accumulation of dominance-effect-based heterosis has been limited since the proposal of GCA-based approaches and very few strategies to exploit it better have been proposed. *We propose the use of a new surrogate of genetic complementation between genetic pools to increase accumulation of dominance effects and heterosis*. We simulated breeding programs to show how reciprocal recurrent selection by genetic complementation would build the dominance-based heterosis but cheaper than GCA-based approaches and used real phenotypic data from hybrid maize to demonstrate the underlying concepts. We found reciprocal recurrent selection by genetic complementation to be an attractive and viable strategy to exploit dominance, build *de novo* heterotic pools and boost the current GCA-based approaches. If demonstrated in practice, we hypothesized that this approach would lower the cost of breeding drastically and contribute to food security.

**Key message:** Heterotic patterns can be developed quickly through genetic complementation surrogates to produce high-performance hybrids at a low cost in diploid species displaying dominance and boost GCA-based approaches in hybrid breeding.

## Introduction

When Shull (1952) referred to the term heterosis as “increased vigor, size, fruitfulness, speed of development, resistance to disease and to insect pests, or to climatic rigors of any kind, manifested by crossbred organisms as compared with corresponding inbreds, as the specific results of unlikeness in the constitutions of the uniting parental gametes” he mostly focused on the positive phenotypic effects but a clear genetic definition was not provided. The difference between crossbred organisms compared with corresponding inbreds occurs because inbreds do not leverage from the dominance interactions, whereas hybrid or non-inbred organisms exploit the immediate advantage of dominance interactions and epistasis. Single-locus dominance is the phenomena where a heterozygote individual tends to reflect more the phenotype of one of the homozygote individuals, and in polygenic traits these multiple dominance interactions can add to a substantial portion of the phenotypic value (Falconer & Mackay, 1996; Crow, 1999). Throughout this paper, we assume dominance (defined as the phenomenon of one variant (allele) of a gene on a chromosome tending to mask or override the effect of a different variant of the same gene on the other copy of the chromosome) as the main contributor to heterosis. When considered at the individual-level, heterosis is referred as mid-parent or best-parent heterosis (difference between the hybrid and the average of both parents, or the best parent respectively) (Falconer & Mackay, 1996), whereas when considered at the population-level heterosis is referred as baseline (difference between the inbred and non-inbred population) and panmictic heterosis (difference between the interpopulation hybrids and the non-inbred populations from each subpopulation) (Lamkey & Edwards, 1999). In the absence of epistasis, the presence of a positive average difference in performance between hybrids and parents implies the presence of directional dominance which led past breeders to develop approaches to exploit it for commercial settings (Hallauer et al., 2010; Lamkey and Edwards, 1999; Birchler et al., 2010). There is overwhelming evidence that heterosis is a manifestation of dominance effects, as opposed to overdominance (see the review by Bingham, 1998). In addition, evidence indicates that heterosis is the opposite effect of inbreeding in which recessive-lethal alleles are unmasked in the phenotype in the absence of epistasis (Davenport, 1908; East, 1936; Falconer and Mackay, 1996; Lamkey and Edwards, 1999; Bernardo, 2002; Varona et al., 2009; Charlesworth & Willis, 2009; Joshi et al., 2015). Since heterosis is the reflect of the accumulation of many genes, dominance x dominance epistatic interactions can contribute but will not be the focus of this paper (Lippman & Zamir, 2007; Jiang et al., 2017).

The idea that genes can have different modes of action − that is, an additive, a dominance, or an epistatic effect on the final phenotype − led scientists like Comstock and Robinson, among others, to develop sophisticated mating designs to understand the gene action of important and complex traits such as grain yield in maize (Comstock & Robinson 1949, 1952; Jinks & Jones, 1957; Bernardo, 2002). Better understanding of the inheritance theory, led to the development of recurrent and reciprocal recurrent selection (RRS) as the predominant method to improve populations, which over the years have been enhanced by additional mating designs and introgression steps to develop products capable of increasing the performance in the agricultural fields in the 20^th^ and 21^st^ century (Hallauer et al., 2010). Although the Green Revolution had an enormous impact in developing countries using major genes (e.g., dwarfing genes) (Evenson & Gollin, 2003; Hedden, 2003), hybrid breeding based on the quantitative genetics theory of dominance was one of the main drivers together with improved agronomic management of the massive yield increases in developed countries in North America and Europe in crops like maize (Hill, 2010). The reciprocal recurrent selection based on recycling parents using GCA as a surrogate of value and SCA as the additional value to identify products, have become the foundation of hybrid breeding until the present (Hallauer et al., 2010). Unfortunately, RRS compared to single-pool inbred breeding (i.e., line breeding) and single-pool non-inbred breeding (e.g., clonally propagated crops) tends to be more expensive in terms of time and money due to the need of additional crossing among subpopulations and evaluation of the resulting hybrids depending on the mating strategy (e.g., testcrossing). In summary, since the proposal of GCA-based approaches, few strategies to harness dominance and epistasis addressing time and resources constraints of GCA-based approaches have been proposed (de Boer & Hoeschele, 1993; Mrode, 2014; Hallauer et al., 2010; Werner et al., 2020).

Here, we want to highlight that the availability of genomic information in the form of genetic markers and our knowledge of the modes of inheritance (especially directional dominance) provides us with a unique opportunity to breed for the accumulation of dominance effects to exploit heterosis in a fast and cheap way to boost RRS approaches that improve populations based on GCA. We have developed a method, a new variant of reciprocal recurrent selection, which we call breeding dominance by genetic complementation which leverages from the idea that dominance alleles mask deleterious alleles to create pools that highly complement each other and display high levels of dominance-based heterosis in a controlled manner. This can be thought as a controlled genetic distance method. We show with simulations and some real datasets the implementation of the method and the implications. This could drastically reduce the cost of a breeding program depending on the levels of dominance found in the species of interest and serve to create *de novo* heterotic pools fast and cheap.

## Materials and Methods

### The complementation approach and computation of surrogates

Under the complete dominance-based heterosis hypothesis, the best individuals are those which accumulate A) all heterozygote interactions or B) all homozygote interaction for the positive allele, genome-wide for the QTLs underlying a trait (e.g., yield). In a diploid species, that implies accumulating numerous genotypes of the form A_i_A_j_ for multiple QTLs (i ≠ j) or A_i_A_i_ (considering A_i_ the positive allele). Selection of individuals for having the maximum performance under complete dominance for a trait under the control of “n” QTLs is given by the probability of 3^n^/4^n^ (Figure 1). Under complete dominance, that is a probability of finding one individual per million individuals for 50 QTLs. It is accepted that for complex traits there are hundreds if not thousands of underling QTLs with varying levels of dominance, from no dominance (fully additive) to full dominance. Assuming -only for demonstration purposes-complete dominance, the only practical way to achieve a fully heterozygote individual for hundreds and thousands of QTLs is to create two selection streams, one that selects for fixation the allele A_i_ in population 1 and another that selects for allele A_j_ in population 2 (where i ≠ j). Although it will require multiple cycles (recurrent selection) to achieve, this remains more affordable than cultivating billions of individuals and attempting to phenotype them accurately to identify the best individual.

**Figure 1.**
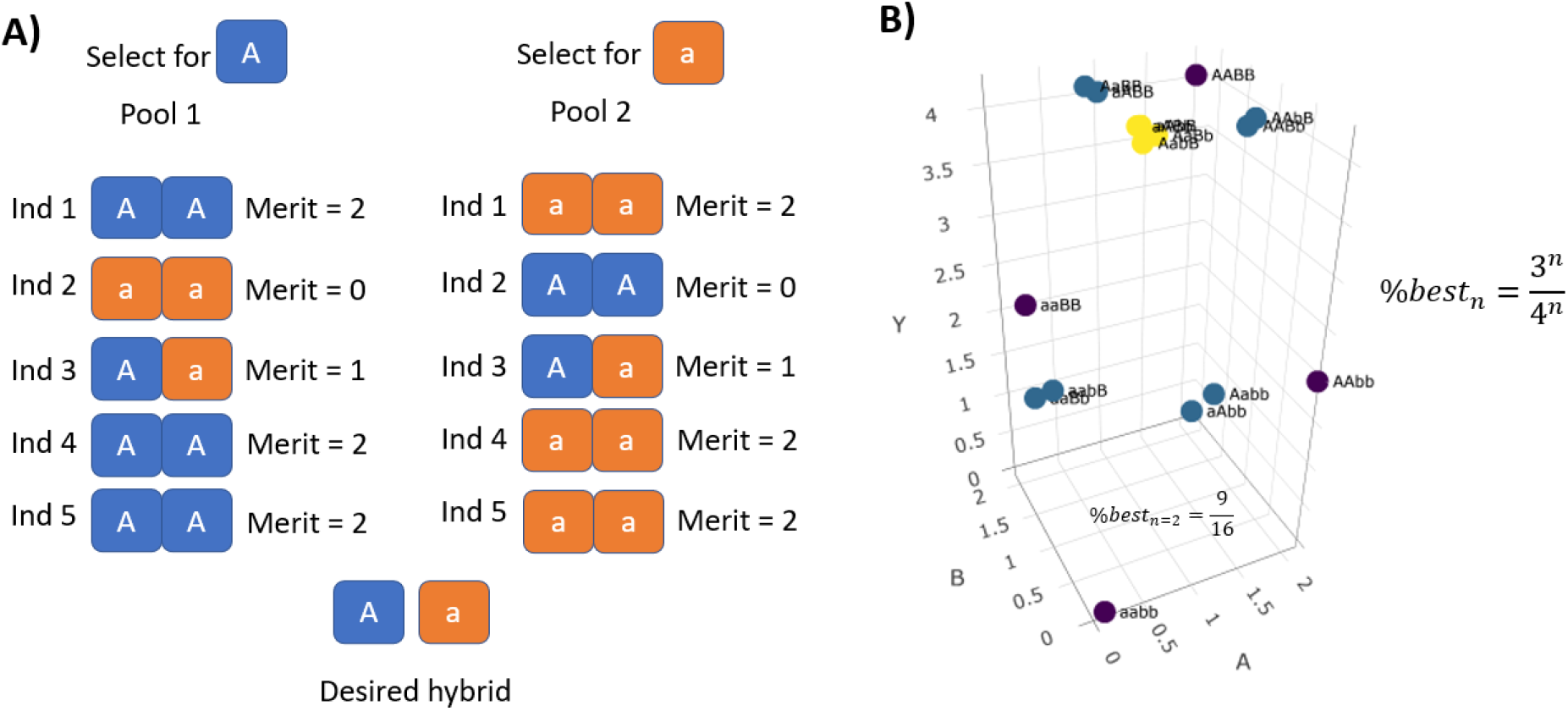
Graphical description of the principles behind reciprocal recurrent selection by genetic complementation. In A) Pool 1 assigns more merit to the selection of one allele whereas the other pool selects for the opposite allele. Both pools are bred to fix the desired allele (or haplotype if reasoning genome-wide). Under the directional dominance model, over time hybrids between these two populations are expected to better complement each other and produce higher-performing hybrids that display strong heterosis. At complete dominance, the best individual is the one that combines all loci at heterozygote state or all the loci at homozygous state for the positive allele. At lower levels of dominance, the individual that combines all loci at heterozygote state is not as good as the individual comprising all the loci at homozygous state for the positive allele. In B) the genetic value (Y) as a function of two QTLs (A and B) is shown, displaying the relationship to find the best hybrids depending on the number of QTLs under complete dominance (3^n^/4^n^).

Following the complete dominance assumption for now (we know that complete dominance does not hold for all loci and different level of dominance exist across the genome), to ensure the creation of that idealized hybrid, we need to ensure the definition of the idealized or desired genotype and haplotype for each of the two subpopulation/pools (in case of diploids). The foundational step of a reciprocal recurrent selection approach by genetic complementation is that two populations will be created with the specific purpose to complement each other (Figure 1).

We then need to define first a desired/idealized genotype under the idea of genomic complementation and complete-dominance assumptions (to be a haplotype in its final state) as the accumulation of homozygous allelic states for all the QTLs behind a trait of interest in a population, and to be opposite/complementary to another population. Notice the statement, under the complete dominance assumption. There are two challenges to implement this: 1) unknown location of the QTLs behind the trait of interest (ideally, we would only focus on the actual QTLs), and 2) unknown coupling-repulsion phases present in the populations (this could potentially slow down the recurrent selection using complementation). To face these challenges, we propose that under the infinitesimal model of complex traits, the use of a genetic-marker chip with thousands of genetic markers is enough to target most QTLs of interest by linkage disequilibrium, and by identifying the highest-frequency alleles in a population (using genetic markers) we can identify what are the alleles in coupling phase or genome-wide population haplotype in order to come up with the desired genotypes for each pool (instead of picking desired alleles at random, which may lead to repulsion-phase linkages in both populations that are difficult to break). Assuming a population of individuals genotyped with biallelic markers coded as 0, 1, and 2 (corresponding to A_i_A_j_, A_j_A_j_) with frequencies (f_1_, f_2_, f_3_), the desired/idealized genotype for the i^th^ marker in the pool 1 is:

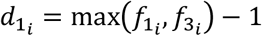

Over the entire genome, the vector d_1_ of desired alleles for pool 1 is all d_1,i_’s for all markers. The vector d_2_ of desired alleles for pool 2 and complementary to pool 1 is just 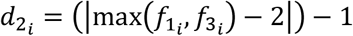. These vectors represent the ideal genotype for all individuals in their respective pool.

Once the desired/idealized genotypes in the vectors d_1_ and d_2_ have been defined, the next step is to define the surrogate of complementation. This will reflect how close a genotype is to the desired/idealized genotype to complement the opposite pool, and is calculated as follows:

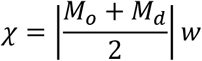

Where M_o_ is the matrix of observed genotypes coded as -1, 0 and 1 for the presence of a reference allele, M_d_ is the matrix of desired genotypes composed by the d vector copied and row-bound as many times as individuals (rows) in the M_o_ matrix. The vector w refers to the weights to be applied to each marker in case we would like to weight by allele frequencies and | is the entry-wise absolute value operator. The vector denoting the complementation value for each individual to the opposite pool is named χ. The higher the value of χ the better parent the individual will be for the next generation to complement the opposite pool moving forward.

Although the complete dominance assumption is unrealistic, for any genome-wide average value of dominance degree between 0 and 1, this approach can help harness that level of heterosis. Additional theoretical details of the complementation approach can be found in Annex 1.

### Validating the complementation model in a generic simulation and real datasets

Under the proposed genetic model, a collateral effect on hybrid performance based on levels of dominance is that dominance effects can be predicted with reasonable accuracy by the complementation surrogate (χ) in different forms (Bernardo, 1992, 2002). We selected parental lines using two variations of the χ metric to predict the hybrid performance in the current generations and keeping track of the correlation between the modified χ metric and the hybrid performance:

1. The complementation of individual i from population A to the desired haplotype of population B (implying only one side contributes to the prediction, the other side is constant for all individuals).
2. The complementation between individuals i and j from pools A and B respectively (implying that both sides contribute to the prediction).

We kept track of the correlation of hybrid performance and the complementation surrogate at different levels of dominance and levels of genetic variance across 10 cycles. The appropriate correlation is expected to be calculated between the dominance effect and the complementation surrogate, but the known difficulties to separate dominance from additive effects motivated us to use the total hybrid performance instead of trying to calculate/separate only the dominance effects which is expected to lower the expected correlation.

Then, we took the hybrid-maize dataset from Kadam et al. (2016) which includes marker data and yield performance for 312 hybrids coming from two heterotic pools (46 lines in the female pool and 172 lines in the male pool) tested in five environments and calculated the one-to-one complementation metrics χ_ii_ and χ_ip_ (individual-to-individual and individual-to-population respectively), calculating the correlation between these metrics with the hybrid yield performance in each of the 5 environments. In addition, we took the maize dataset made available by Technow et al. (2014) that includes genetic marker data and adjusted means across environments data for 1254 hybrids coming from crosses between the flint (86 lines) and dent (123 lines) maize pools and 35,478 SNP markers, calculating the one-to-one complementation metrics χ_ii_ and χ_ip_ and the correlation between these metrics with the hybrid yield performance across environments.

### Simulating the influence of different levels of dominance and number of QTLs in building heterosis

#### Baseline

As a baseline, we simulated a generic hybrid breeding program with a three-stage evaluation strategy, in addition to the crossing block stage and four steps of single seed descent where planting material is multiplied, and inbred lines are developed. These evaluation stages include stage 1 (*per se* evaluation), stage 2 (first testcross with 3 testers) and stage 3 (topcross). The pipeline began with 100 parental lines in each of the two simulated pools. From the 100 parents, 50 crosses were made, each with 20 progeny, thus resulting in 1000 individuals. All 1000 went through single seed descent (SSD) for 4 generations and then evaluated at stage 1 in one environment and one replication per environment. At the same time as *per se* evaluation occurs (only used for advancement, not recycling), the testcrosses of the 1000 lines are performed against 3 testers of the opposite pool. In Stage 2, the testcrosses are evaluated in two environments and two replications per environment. From stage 2 evaluation, the best 20 families and best 5 lines per family in each pool were selected and advanced to stage 3 where 100 hybrids are made and tested in two environments and three replications per environment. Recycling of parents occurred after the evaluation of Stage 2 when the best 100 lines in each pool are selected to become parents of the next generation (6^th^ year). We were interested in understanding if the complementation approach is effective to build heterosis compared to classical recurrent and reciprocal recurrent selection approaches. The availability of a SNP chip with 500 markers was assumed at any stage that required genotyping. The baseline treatment was named TWO_POOL_GCA_PHENO.

#### Alternative treatments

To keep the treatments comparable we used the same number of individuals, the only difference between treatments is the time when recycling of parental lines occurs and the surrogate of merit used. The scheme that only uses complementation surrogate to recycle (TWO_POOL_COMP) is expected to recycle after 1 year. A positive control treatment recycling based on the true GCA in the 6^th^ year (TWO_POOL_GCA_TRUE) follows the same formula according to Schnell (1965) and Rembe et al. (2019):

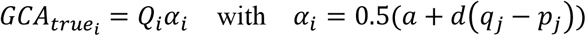

Where i and j refer to the pools i and j, α is the vector of average effects of an allelic substitution, a and d are the vectors of additive and dominance effects of the QTLs respectively, p and q are the vectors of allele frequencies, and Q is the matrix of QTL calls coded as 0, 1 and 2 for the i^th^ pool.

The negative controls included selecting parents at random in Stage 2 in the 6^th^ year (NEGATIVE_CONTROL), and selecting parents based on breeding value in two independent pools to make hybrids as final product (TWO_POOL_BV; we did use the genetic value but in an inbred line this is expected to be equal to the breeding value in the absence of epistasis), which is not expected to build dominance-based heterosis (panmictic) since it is focused on the additive portion of the total genetic value (only builds some panmictic heterosis initially due to the initial drift but does not keep building it long term). In addition, a treatment where a program runs the complementation approach for 5 years prior to switches to formal RRS was included (TWO_POOL_COMP_TO_GCA). As such, a total of 6 simulation treatments were defined (Figure 2). To identify the best strategy to build heterosis, the stochastic genetic simulation was conducted in the R package AlphaSimR (Gaynor et al., 2021).

**Figure 2.**
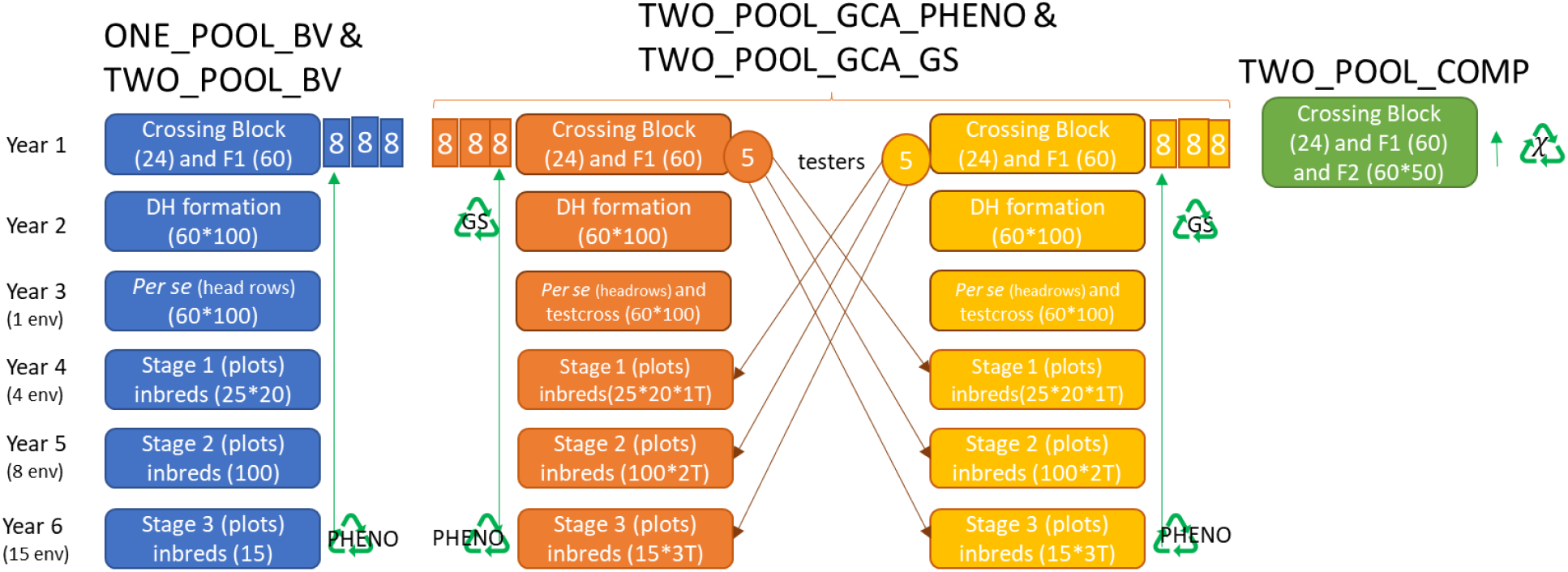
Summary of treatments (breeding strategies) compared to build dominance-based heterosis. Each treatment uses a different surrogate of merit to select parents in each pool.

#### Burn-in genome sequence

For each replicate, a genome consisting of 1 chromosome was simulated for the hypothetical plant species. These chromosomes were assigned a genetic length of 1.43 Morgans and a physical length of 8 × 10^8^ base pairs. Sequences for each chromosome were generated using the Markovian coalescent simulator (MaCS; Chen et al., 2009) implemented in AlphaSimR (Gaynor et al., 2021). Recombination rate was inferred from genome size (i.e., 1.43 Morgans/ 8 × 10^8^ base pairs = 1.8 × 10^−9^ per base pair), and mutation rate was set to 2 × 10^−9^ per base pair. Effective population size was set to 30 to mimic an evolution history of natural and artificial selection.

#### Burn-in founder genotypes

Simulated genome sequences were used to produce 100 founder inbred individuals. These founder individuals served as the initial parents in the burn-in phase. Sites segregating (2000) in the founders’ sequences were randomly selected to serve at different number of quantitative trait nucleotides (QTN) per chromosome for each of the defined treatments: 1, 10, 50, 100, 1000.

#### Burn-in phenotypes

A single highly complex trait for the different number of QTNs specified above was simulated for all founders. The genetic value of this trait was determined by summing its QTN allelic effects. The dominance effect (*d*) at a locus is then its dominance degree (*δ*) times the absolute value of its additive effect, with scaling to achieve user-specified additive or total genetic variance. Dominance effects must be scaled even if use of additive variance is specified (useVarA = TRUE) because dominance effects contribute to additive variance.

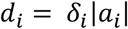

At the same time, the allele effects followed this formula:

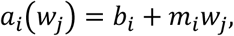

where *a*_*i*_ is the allele effect for QTN *i, w*_*j*_ is the environmental effect for year *j, b*_*i*_ is the QTN intercept and *m*_*i*_ is the QTN slope on the environmental effect. The slope, intercept, and environmental effects were sampled from the following Gaussian normal distributions. Details on the full formulation of genotype by environment interaction simulation features enabled in AlphaSimR can be found in Gaynor (2021), and is implemented in the addTraitAD() function through the mean dominance degree (meanDD) argument. The values set for the mean dominance degree were 0 (purely additive), 0.5 (partial dominance), and 0.95 (almost complete dominance). The varDD argument was set to 0.2.

The genetic values of each inbred individual were used to produce phenotypic values by adding random noise sampled from a Gaussian normal distribution with mean 0. The variance of the random error was varied according to the three stages of evaluation defined in the breeding program to achieve the following heritabilities: STG1 H^2^=0.2, STG2 H^2^=0.5, STG3 H^2^=0.5. All this was achieved with the setPheno() and setPhenoGCA() functions and the varE argument. A summary of simulation features for the genome and phenotypes can be found in Table 1.

**Table 1.** Summary of simulation features for the genome and phenotypes.

| Simulation features |  |  |  |
| --- | --- | --- | --- |
| Burn-in |  | Generic genetic model | Inbred and inbred-hybrid programs |
|  | Genome sequence | 100,000 generations of evolution | 100,000 generations of evolution |
|  |  | 1 chromosome pair | 18 chromosome pairs |
|  |  | 1.43 Morgans per Chromosome | 1.43 Morgans per Chromosome |
|  |  | 8 x 10<sup>8</sup> base pairs per chromosome | 8 x 10<sup>8</sup> base pairs per chromosome |
|  |  | 2 x 10<sup>-9</sup> mutation rate | 2 x 10<sup>-9</sup> mutation rate |
|  | Founder genotypes | 100 inbred founders per pool | 24 inbred founders per pool |
|  |  | 1, 10, 50, 100, 1000 QTNs (additive+ dominance effects) | 1000 QTNs (additive+ dominance effects) |
|  |  | Normally distributed QTN effects with meanDD values of 0, 0.5 and 0.95 and varDD=0.2 | Normally distributed QTN effects with meanDD value of 0.5 and varDD=0.2 |
|  |  | 0 cycles of breeding | 20 years of breeding |
|  |  | Inbred individuals | Inbred individuals |
|  |  | Conventional breeding | Conventional line and hybrid breeding |
| Evaluation | Future breeding | 20 cycles of breeding | 40 years of breeding |
|  |  | Testing alternative allocation of resources | Testing alternative allocation of resources |
|  |  | Equal cost programs | Variable cost programs |
|  |  | Conventional breeding and alternative treatments | Conventional breeding and alternative treatments |

Population means and standard errors for additive, dominance and total value at Stage 2 of yield evaluation across 20 cycles of selection for the treatments described previously were computed using the dplyr library in R (Wickham et al., 2021), and plotted using the ggplot2 library in R (Wickham, 2011). Twenty replicates were run for each simulation treatment.

#### Simulating the effect of updating testers to build the dominance

Using the previous simulation and the baseline treatment of reciprocal recurrent selection based on GCA (TWO_POOL_GCA_PHENO), we tested different strategies to update the testers. We simulated the following treatments: RRS without updating the testers (TWO_POOL_GCA_UPDATE0), and RRS updating all the testers every 1, 4, and 8 cycles (TWO_POOL_GCA_UPDATE_x, with x={1, 4, 8}), and as control treatments: RRS using true GCA updating all the testers every cycle (TWO_POOL_GCA_TRUE (positive control), and RRS selecting parents at random (NEGATIVE CONTROL). Four levels of dominance were considered (meanDD equal to 0, 0.2, 0.5, and 0.9). Additive, dominance and total genetic value were recorded as in the previous simulation.

### Simulating the implementation of the complementation approach in a running inbred program

To understand how a program that currently improves inbred lines would transition into a hybrid program under either the conventional strategy recycling based on GCA or the proposed complementation strategy to build initial pools and then move to recycle based on GCA, we simulated the same treatments as before but for the genome structure of a crop similar to maize, and a strategy that reflects how a program working on recycling and delivering inbreds would transition to the complementation approach.

#### Baseline

As a baseline scheme, we simulated a generic inbred breeding program with a four-stage evaluation strategy in addition to the crossing block stage and a double haploid step where planting material is multiplied and inbred lines are developed. The evaluation stages include Stage 1 (*per se* evaluation of short rows in one location in a single rep), Stage 2 (big plots single rep at 4 locations), Stage 3 (big plots single rep at 8 locations), Stage 4 (big plots single rep at 15 locations). The pipeline began with 24 parental lines in each of the two simulated pools. From the 24 parents, 60 crosses were made, each generating 130 double haploid (DH) lines, thus resulting in 16,900 DHs. From the 16,900 lines evaluated in Stage 1, the best 25 families and the best 40 DHs per family were selected (1000 lines). In Stage 2 the best 100 lines across all families are selected. In Stage 3, the best 15 lines are selected for on-farm trials. Recycling of parents occurred after the evaluation of Stage 3 where the best 8 out of the best 15 parental lines replace 8 lines of the 24 elite parents. Since we were interested in understanding if the complementation approach can be implemented effectively to build heterosis compared to adopting immediately reciprocal recurrent selection that uses GCA to recycle parents, we used this program as the baseline (burn-in) for the first 20 years for all the next treatments. This scenario was named two pool selection based on breeding value (TWO_POOL_BV) since we assumed two programs running in parallel to simulate interpopulation hybrids that serve as control since selection based on BV (instead of GCA) in opposite pools is not expected to build dominance (Hallauer et al., 2010). A SNP chip with 5000 markers was assumed available at any stage of the program that required genotyping.

#### Alternative treatments

To transition the inbred program into a hybrid program, we simulated a treatment in which, following 20 years of burn in, classical reciprocal recurrent selection took place through the use of testcrosses to generate hybrids that can be evaluated to calculate GCA that can be used for recycling parents. Following the evaluation of 16,900 DHs in short rows the best 20 DHs from the best 25 families (500 lines) were testcrossed against the best elite (tester) from the opposite pool (Stage 2), then the best 100 lines against the 3 best lines (testers) of the opposite pool (Stage 3), and finally the best 15 lines against the best 5 lines (testers) of the opposite pool (Stage 4). Recycling of parents occurred after the evaluation of Stage 3 where the best 8 out of the best 15 parental lines replace 8 lines of the 24 elite parents but since this is based on GCA it took one season later compared to the convention line approach. This scenario was named reciprocal recurrent selection based on GCA (TWO_POOL_GCA_PHENO). After Stage 3, 4 of out the 5 testers are replaced with the top selected lines from Stage 3 (Figure 2). A similar reciprocal recurrent selection treatment based on GCA but using genomic selection (a training population based on hybrid phenotypes) to predict the GCA of DH lines early in each pool and recycle much faster was also tested (TWO_POOL_GCA_GS).

For the complementation approach, we used a similar hybrid program to conventional RRS as described above, but recycling was based on the complementation surrogate against the opposite pool using the χ metric in a portion of the DHs available [a random sample of 3000 was assumed to be genotyped, the χ metric was calculated, and we selected the best individual within each family)], directly reducing the cycle time since there is no need for extensive phenotypic evaluation of individuals for complex traits. Still, this scenario assumes only 2 cycles per year. This scenario is named reciprocal recurrent selection based on genetic complementation (TWO_POOL_COMP). A variation of this approach is the one that combines complementation and GCA where complementation is applied for twelve years and then GCA takes off, named reciprocal recurrent selection based on genetic complementation and GCA (TWO_POOL_COMP_TO_GCA). All RRS approaches (GCA and complementation) formed the initial heterotic pools by splitting a sample of DH lines randomly into two *de novo* subpopulations (Figure 2).

The negative control was based on selecting parents at random in Stage 3 (NEGATIVE_CONTROL). Positive control were selection based on true GCA (TWO_POOL_GCA_TRUE) and selection based on true genetic complementation (TWO_POOL_COMP_TRUE) where QTLs are assumed to be known (only as control, the complementation approach doesn’t require QTLs to be known). As such, a total of 4 simulation treatments and 3 controls were defined. To quantify the effect of implementing these strategies to build heterosis, a stochastic genetic simulation was conducted in the R package AlphaSimR (Gaynor et al., 2021). These set of simulations considered differences in cycle time between the different strategies.

#### Burn-in genome sequence

For each replicate, a genome consisting of 10 chromosome pairs was simulated for the hypothetical plant species similar to maize using the argument species=“MAIZE” in the runMacs() function available in AlphaSimR that simulates the species evolutionary history of maize. These chromosomes were assigned a genetic length of 1.43 Morgans and a physical length of 8 × 10^8^ base pairs. Sequences for each chromosome were generated using the Markovian coalescent simulator (MaCS; Chen et al., 2009) implemented in AlphaSimR (Gaynor et al., 2021).

#### Burn-in founder genotypes

Simulated genome sequences were used to produce 24 founder inbred individuals. These founder individuals served as the initial parents in the burn-in phase. Sites segregating (800) in the founders’ sequences were randomly selected to assign 300 quantitative trait nucleotides (QTN).

#### Burn-in phenotypes

A single highly complex trait for the 3000 QTNs specified above was simulated for all founders. The genetic value of this trait was determined by summing its QTN allelic effects having an additive and a dominance value as mentioned in the section for the generic simulation.

## Statistical analysis

Stochastic simulations executed with AlphaSimR included twenty Monte-Carlo replications. For the exploration of the practical implementation of genetic complementation in inbred and hybrid programs, standard errors (SE) were computed for treatments across years as:

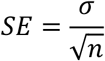

Where σ is the standard deviation from the twenty Monte-Carlo replicates and n is the number of replicates (20). Standard errors were plotted as shadowed lines for the genetic trends in all figures to declare differences between treatments.

## Results

### Validating the complementation model in a generic simulation and real datasets

Given the unavailability of data from programs that have executed the complementation approach for several cycles as proposed, we had to incorporate a set of the side results from the complementation approach. Under the proposed genetic model, a collateral effect is that the hybrid performance can also be predicted with more or less accuracy by the complementation surrogate (χ) depending on the levels of dominance (Table 2). In theory, the complementation surrogate (χ) is highly correlated with the dominance effect observed in hybrids, but we had to focus hybrid performance given the difficulty to estimate properly or orthogonally the dominance effects (Nishio and Satoh, 2014).

**Table 2.** Genetic correlation and VarD/VarG between the complementation surrogate χ_ip_ (individual-to-population complementation) and total genetic (additive + dominance) value of the hybrid performance at different levels of dominance degree in a simulated program. Genetic correlation is shown in cells and VarD/VarG is shown inside parenthesis.

| **meanDD/Cycle** | **1** | **4** | **7** | **10** |
| --- | --- | --- | --- | --- |
| 0 | -0.06 (0.12) | -0.03 (0.18) | -0.07 (0.14) | 0.18 (0.26) |
| 0.5 | 0.62 (0.84) | 0.58 (0.55) | 0.43 (0.43) | 0.06 (0.4) |
| 0.95 | 0.77 (0.89) | 0.83 (0.86) | 0.44 (0.81) | 0.3 (0.7) |

We kept track of the correlation of hybrid performance and the complementation surrogates at different levels of dominance and levels of genetic variance across 10 cycles in the simulation of a generic breeding program (Table 2). The χ metric is a value for the line merit instead of hybrid performance, so to produce a χ metric for the hybrids themselves we developed the individual-to-population χ_ip_ and individual-to-individual χ_ii_ metrics for a cross. We found that at intermediate and high levels of dominance (meanDD of 0.5 and 0.95 respectively) the correlation between the complementation surrogate and the total-genetic value of the hybrids were intermediate to high (Table 2). At no dominance (meanDD=0 or purely additive trait) the correlations were around zero at any cycle of selection (Table 2). At intermediate levels of dominance (meanDD=0.5) the levels of correlation between hybrid performance and χ_ip_ ranged from 0.4 to 0.6 in the first 7 cycles of selection, decreasing at a constant rate as dominance variance gets depleted (Table 2). For almost complete dominance (meanDD=0.95) the levels of correlation between hybrid performance and χ_ip_ ranged from 0.4 to 0.8 in the first 7 cycles of selection, decreasing at a constant rate as dominance variance is depleted (Table 2). The higher correlation observed is between the complementation surrogate and the dominance effects.

Using the hybrid-maize dataset from Kadam et al. (2016), which includes marker data and yield performance for 312 hybrids coming from two heterotic pools (46 lines in the female pool and 172 lines in the male pool tested in five environments) and calculated the one-to-one complementation metrics χ_ip_ and χ_ii_, and calculated the correlation between these metrics with the hybrid yield performance in each of the 5 environments (Figure 3). We found the correlations between hybrid performance and χ_ii_ (individual-to-individual complementation) to range between -0.2 and 0.1 (Figure 3a), whereas the correlations between hybrid performance and χ_ip_ (individual-to-population complementation) ranged between -0.3 and 0.45, being more consistent and oscillating between positive and negative depending on the average performance for the environment (Figure 3b). Interestingly, the direction of the correlation changed depending on the environment yield. Positive correlations were found in high-performing environments and negative correlations were found in stressful environments. In addition, using the maize dataset made available by Technow et al. (2014), the correlation between hybrid performance and χ_ii_ to be 0.003, whereas the correlation between hybrid performance and χ_ip_ was 0.41 (Supplemental Figure 1).

**Figure 3.**
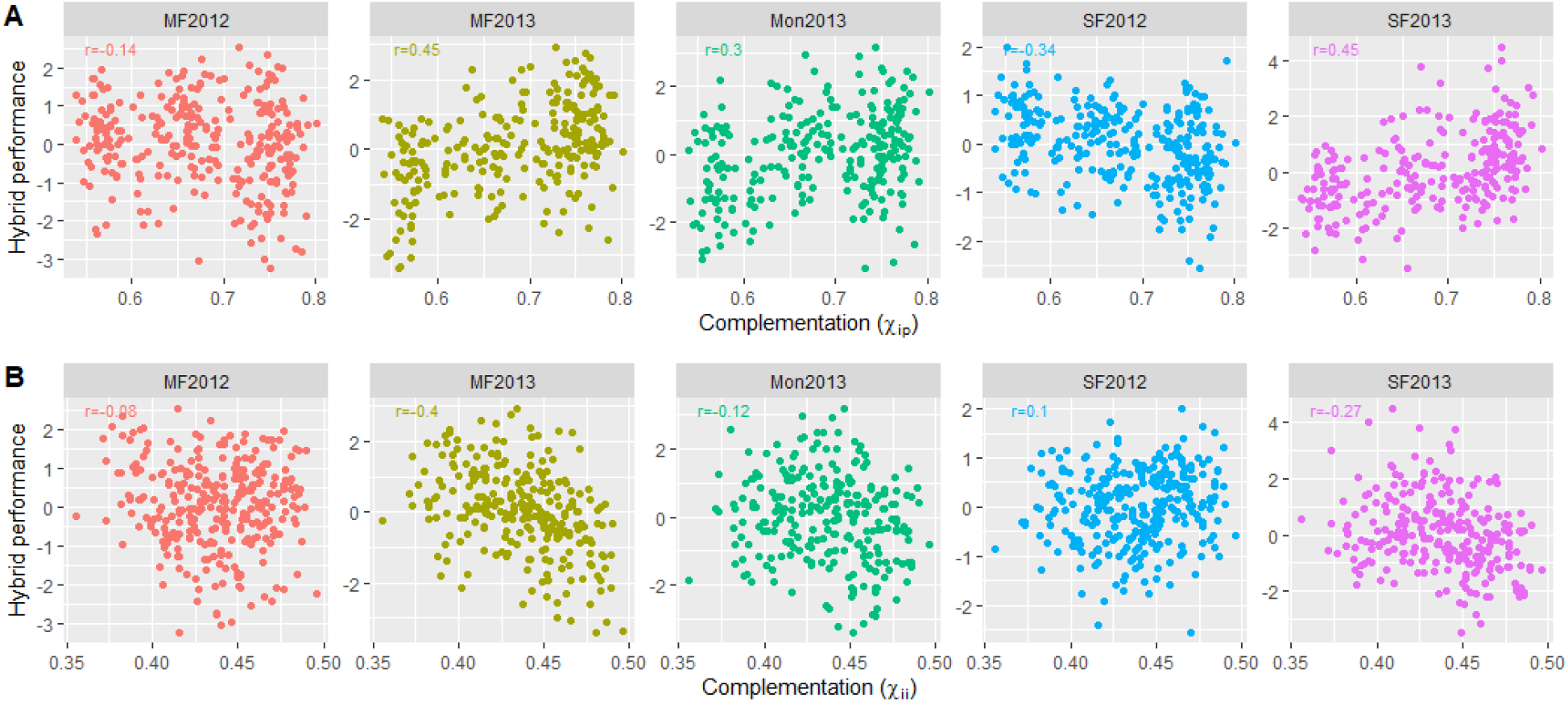
Genetic correlation between the complementation surrogates (individual-to-population and individual-to-individual genetic complementation) and hybrid performance at five different environments. Columns represents the five different environments where hybrids were tested. A) represents the individual-to-population genetic complementation, B) represents the individual-to-individual genetic complementation. Correlation legend is shown in the upper left corner of each plot. Data comes from Kadam et al. (2016) and comprises a real dataset of hybrids between the dent and flint heterotic pools.

### The influence of dominance degree and number of QTLs in total genetic value

To increase our understanding of how the complementation χ metric can increase genetic gain for dominance, we expanded the simulation not only to different levels of dominance (no dominance to complete dominance), but also to different trait complexities (i.e., varying levels of quantitative trait nucleotides [QTLs] behind the trait) and compared it to the classical GCA approach. When looking exclusively at dominance gain we found that, independently of the number of QTLs behind the trait, the complementation (TWO_POOL_COMP & TWO_POOL_COMP_TO_GCA) and the RRS strategies (TWO_POOL_GCA_PHENO & TWO_POOL_GCA_TRUE) strategies were able to build dominance effects efficiently, whereas the negative control (TWO_POOL_BV) did not increase it (Figure 4, Supplemental Figure 2). At all levels of dominance, the complementation approaches were shown to be as effective to increase the dominance values as selecting based on GCA. On the other hand, when looking at the additive gain, we found the GCA-based and BV-based approaches to be the only strategies able to increase this value (Figure 4, Supplemental Figure 2). The total genetic gain was positive for all strategies ranking first the RRS methods followed by complementation and last single pool RS based in BV when the mean dominance degree was ≥ 0.5. At lower levels of mean dominance degree single pool RS based in BV ranked first followed by RRS and complementation last. In addition, we found that trying to increase the dominance value in the breeding populations is not possible when testers are not updated (Figure 5).

**Figure 4.**
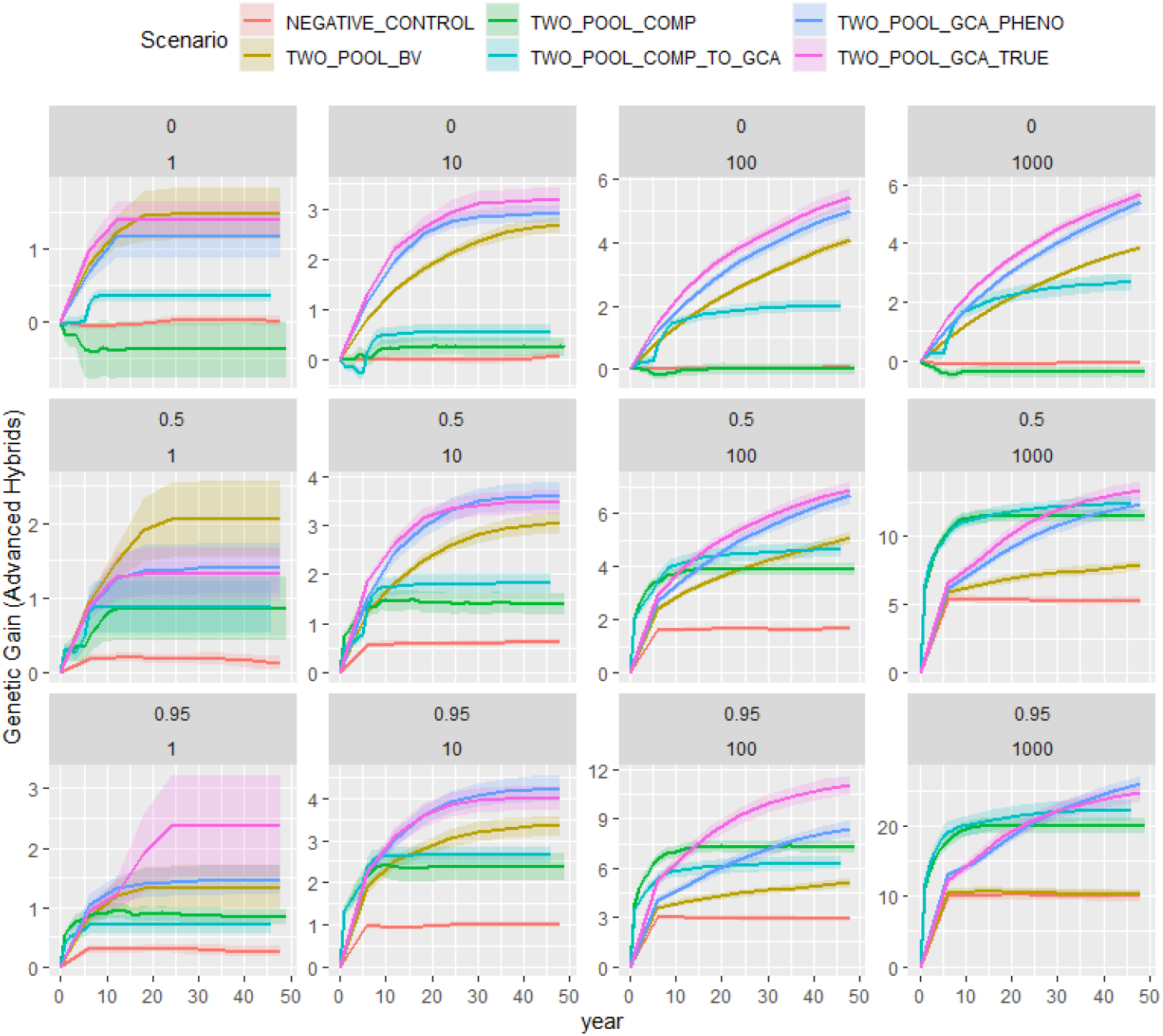
Total genetic gain for a trait in on-farm hybrids under different mean dominance degrees and number of QTLs for different simulated breeding strategies. The columns represent the comparison of strategies where the simulated trait has different number of QTLs behind (1, 10, 50, 100, 1000 QTLs). The rows represent the comparison of strategies where the simulated trait has different levels of mean dominance degree (0 implies a completely additive trait, 0.5 partial dominance, and 0.95 represents a trait with almost complete dominance). A value of varDD=0.2 was used across all scenarios. In the x axis, the number of selection cycles (20) are indicated whereas in the y axis the gain is shown. The different lines represent the different selection strategies: 1) reciprocal recurrent selection program that selects and recycles parents based on GCA (TWO_POOL_GCA_PHENO), 2) reciprocal recurrent selection program that selects and recycles parents based on genetic complementation (TWO_POOL_COMP), 3) reciprocal recurrent selection program that selects and recycles parents based on genetic complementation using true QTLs (TWO_POOL_COMP_TRUE; positive control 1), 4) reciprocal recurrent selection program that selects and recycles parents based on true GCA (TWO_POOL_GCA_TRUE; positive control 2), 5) recurrent selection program that selects and recycles parents based on breeding value (TWO_POOL_BV; negative control 1), 6) recurrent selection program that selects and recycles parents at random in a single pool (NEGATIVE_CONTROL 2).

**Figure 5.**
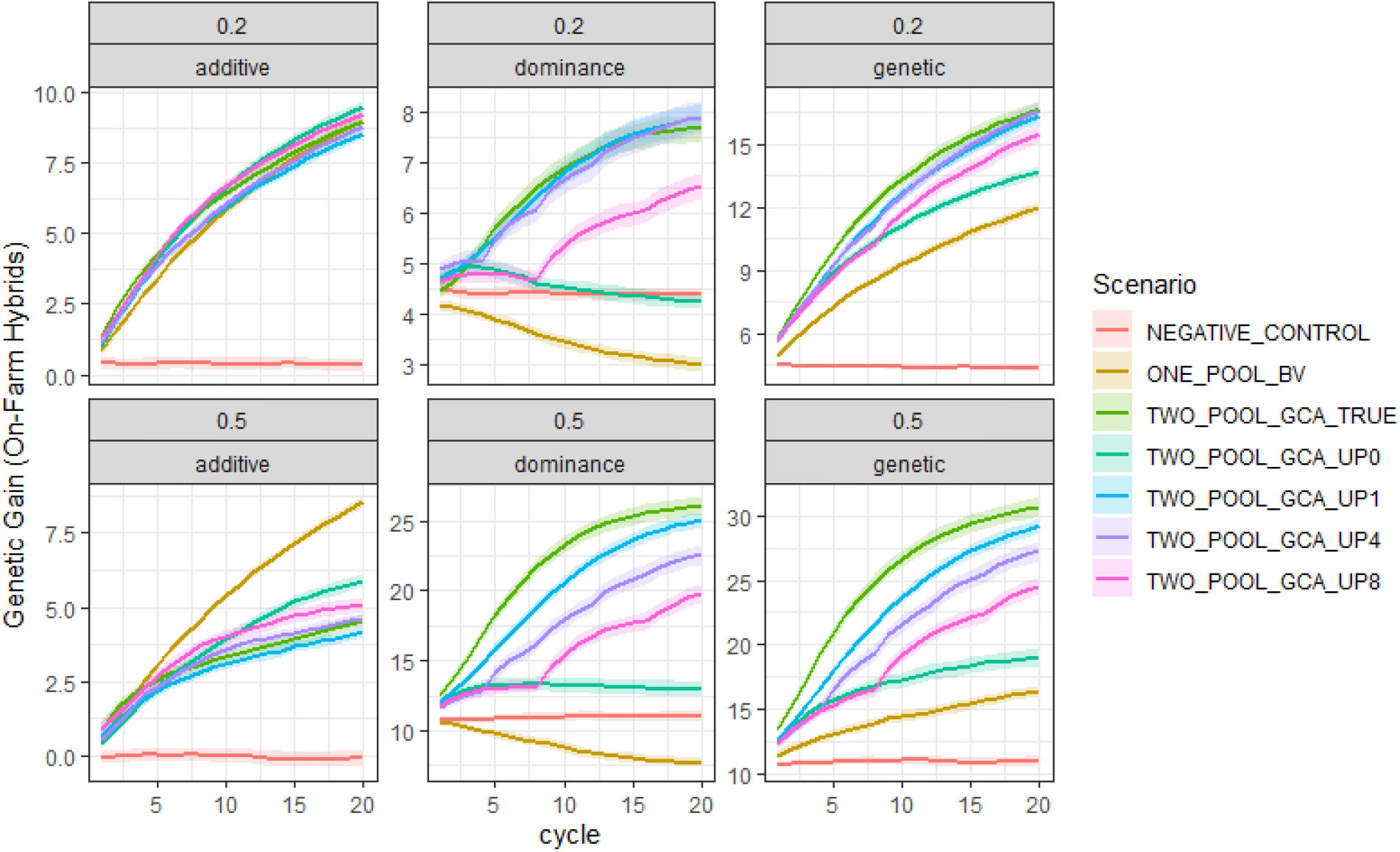
Effect of the representativeness of testers in the increase of dominance-based heterosis. The additive, dominance and total-genetic value increment (columns) based on representativeness of the tester (with respect to its pool) achieved by updating the testers after none, 1, 4, or 8 cycles of selection (treatment lines) at two different levels of dominance degree (rows) is displayed. If the tester(s) are updated often (TWO_POOL_GCA_UP1) they better represent the pool they belong to, and dominance increases at a higher rate compared to programs not updating the testers (TWO_POOL_GCA_UP0) or not that often (rest of the treatments).

### How to implement RRS by genetic complementation in an active inbred program

To understand the best way to implement the genetic complementation approach in an ongoing program that selects inbred materials (i.e., close-to homozygote lines such as rice or wheat), we simulated a breeding program that follows the structure of a line crop. The simulation was initiated with a single population/pool that follows a conventional recurrent selection strategy comprised of a crossing block, a segregation step, a multiplication phase (four generations of single seed descent) and multiple stages of phenotypic evaluation in the target population of environments (TPE) (Figure 6).

**Figure 6.**
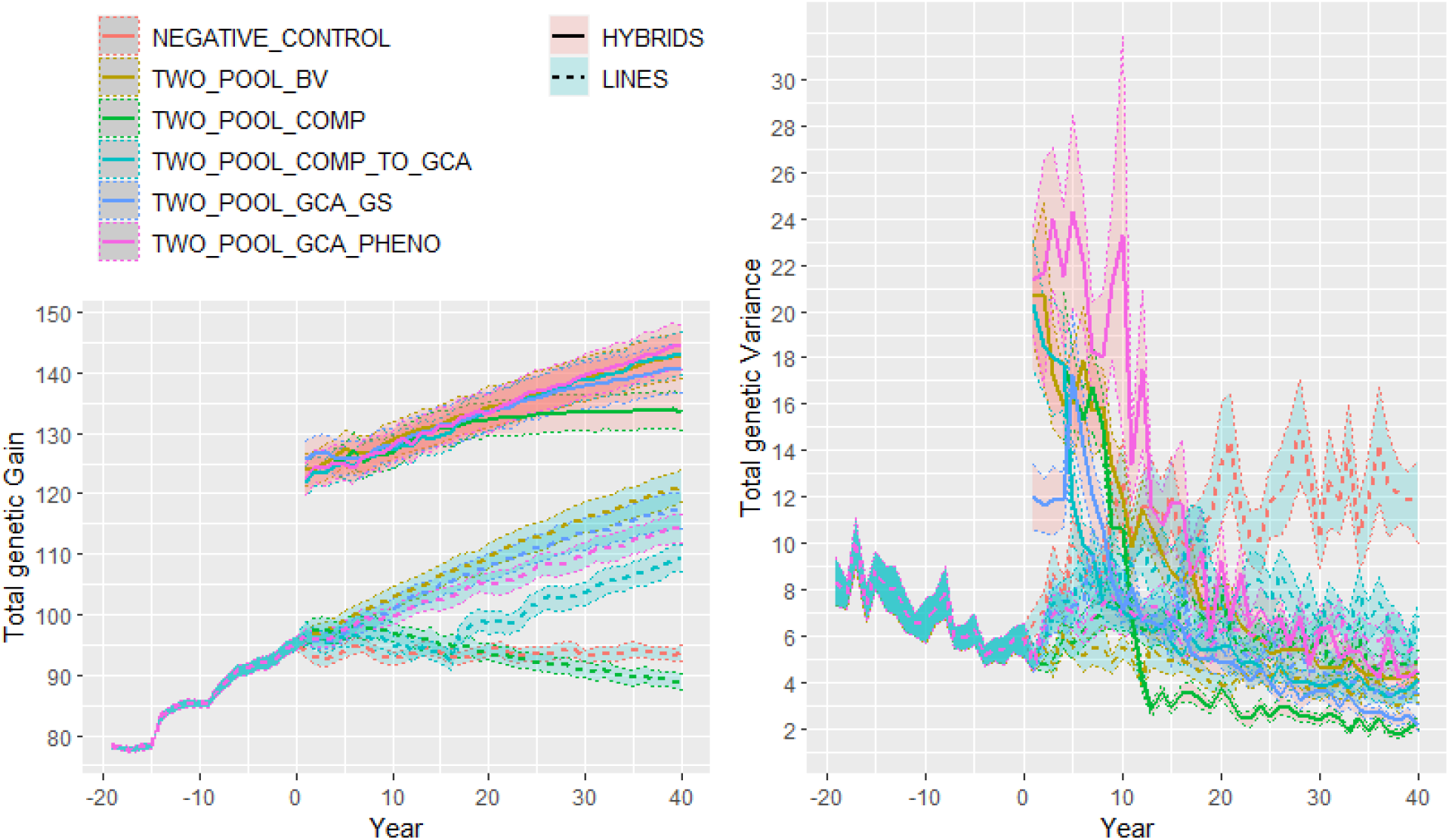
Genetic gain for a complex trait (1000 QTLs behind) with intermediate mean dominance degree (0.5) measured in on-farm hybrids (solid lines) and parental lines (dotted lines) for a simulated line program transitioning to a hybrid program with different breeding strategies. In the x axis, the number of years is indicated whereas in the y axis the genetic gain for total-genetic (A) additive (B) dominance (C) value are shown. The different lines represent the different selection strategies tested: 1) line program continuing the recurrent selection based on breeding value in two independent pools and making hybrids among the pools (TWO_POOL_BV; positive control; pink lines), 2) line program transitioning to RRS to select and recycle parents based on GCA (TWO_POOL_GCA_PHENO; blue lines), 3) line program transitioning to RRS to select purely on complementation (χ) (TWO_POOL_COMP; brown lines, and 4) negative control selecting lines at random (NEGATIVE_CONTROL; red lines). Every year 4 out of the top 5 testers is updated. The value of varDD was 0.2 across all scenarios. Shadow lines represent standard errors for treatments across 20 Monte-Carlo replications.

In the case of the positive control or conventional line program (TWO_POOL_BV) the same rate of response to selection with respect to the burn-in strategy across the 40 years was observed as expected, being that this is the continuation of the same burn-in strategy (Figure 6). The negative control showed no increase in additive, dominance and total genetic value as would be foreseen under random selection. In the case of the program converted to traditional GCA-based RRS (TWO_POOL_GCA_PHENO) we found a lower rate of additive gain but a higher rate of dominance gain resulting in an overall increased rate of total genetic value in the hybrid populations.

The RRS strategy using only the complementation surrogate for selection without phenotyping (TWO_POOL_COMP) was able to produce hybrids with similar performance to the conventional GCA-based RRS approach during the first 15 years after the split due to a substantial increase in dominance and consequently total genetic value (Figure 6, Supplemental Figure 3), but the additive value in the hybrids and lines decreased in performance due to inbreeding (Figure 6, Supplemental Figure 3) causing a stagnation in hybrid performance increase after year 15. Although not shown, using better the genetic variance (e.g., using maximum avoidance or optimal contribution methods) can increase the window of dominance gain. The reciprocal recurrent genomic selection strategy (TWO_POOL_GCA_GS) showed greater gain than its phenotype-based counterpart (TWO_POOL_GCA_PHENO) but did not increase dominance much faster than the complementation approaches. Both, phenotype-based and GS-based RRS strategies can be potentially boosted by the complementation strategy proposed.

## Discussion

### Validating the complementation model in a generic simulation and real datasets

Under complete dominance, the expected correlation between the hybrid performance and the metrics χ_ip_ and χ_ii_ is high since the genetic value is mainly driven by the dominance effects (Table 2), which at the same time are linked to heterozygosity. But as the mean dominance degree decreases the predictive ability of the complementation surrogate decreases (Table 2). The generic simulation model shows that as dominance degree increase the complementation surrogate can be useful for breeding programs to predict the dominance component of the hybrid performance.

The correlation values between the χ_ip_ metric and the hybrid performance found using the dataset from Kadam et al. (2016) and Technow et al. (2014), are closer to the correlation values found in the simulation for intermediate levels of dominance (meanDD∼0.5) which are very similar to the degrees of dominance observed in maize at equilibrium in Nebraska’s and North Carolina’s experiments in the 1980’s and 1990’s (Bingham, 1998; Doebley, 2004; Duvick et al., 2004). Since simulations for the adoption of the complementation approach showed an advantage at intermediate and high levels of dominance, programs currently working as line programs are suitable candidates for the adoption of genetic complementation to increase dominance value and build heterosis (and *de novo* heterotic pools) more efficiently. The methodology does not focus on the initial split of germplasm in pools as the methodology proposed by Zhao et al. (2015), but in the formation of pools through the accumulation of dominance complementations through recurrent selection.

Regarding the low correlation values of χ_ii_ complementation (i.e., individual-to-individual complementation) with the hybrid performance, compared to the χ_ip_ complementation (i.e., the population level complementation), our hypothesis is that since the complementation is defined with respect to a desired haplotype of the opposite population pool, and the complementation at the individual level loses accuracy and relevance.

### The influence of dominance degree and number of QTLs in total genetic value

The fact that RRS strategies (TWO_POOL_GCA_PHENO & TWO_POOL_GCA_TRUE) strategies were able to build dominance effects efficiently, whereas the negative control (TWO_POOL_BV) did not increase it is because the separation of pools at random without a complementation surrogate or a GCA becomes a random process where complementary alleles are not guaranteed to go to the opposite pools and both pools may end up fixing the same allele. Building genetic distance alone is not expected to increase the dominance efficiently. This lack of correlation between genetic distance and heterosis has been observed in several studies in the past (Lamkey et al., 1987; Charcosset, 1991; Bernardo, 1992). Although the genetic complementation is based on the same principle, this can be thought as a controlled genetic distance that guarantees to maximize the heterotic response.

Previously, genetic distance has been proposed as a potential predictor of hybrid performance, with correlation between hybrid performance and genetic distance ranging between 0 and 0.3 (Frei et al., 1986; Bernardo, 1992; Zhang et al., 1996). Bernardo (1992) for example, found that for different values of allele frequencies between set A and set B, the correlation between μ_ij_ (performance) and D_ij_ (distance) was rμ_ij_D_ij_ =0.25, whereas with partial dominance the correlation between μ_ij_ and D_ij_ decreased to ru_ij_D_ij_ =0.13. In other empirical studies, correlations of 0.09, 0.14, 0.32, and 0.46 were obtained by Godshalk et al. (1990), Dudley et al. (1991), Melchinger et al. (1990), and Lee et al. (1989), respectively. This is unsurprising under the directional dominance theory where heterosis is the accumulation of the dominance effects, but not the total performance that includes the additive effects. What builds and maximizes heterosis (dominance value) is the divergence of two pools (for a diploid species) in a controlled manner to put complementary/opposite alleles (maybe even indels) or haplotypes in each pool. Splitting two population/pools without a control builds genetic distance and some heterosis, but permits the same and different alleles (and small-effect mutations) to be increased or fixed in both populations by genetic drift (Charlsworth, 2009). In this case, it is most likely that populations which diverged for opposite alleles are the ones that will display greater heterosis (dominance effects) when crossed back (e.g., flint and dent population in maize), but there should be many others that have diverged in average but which have fixed more similar alleles and therefore show less heterosis (Lande, 1976; Allendorf, 1986; Lynch & Walsh, 1998; Lynch et al, 2016). One of the reasons why we found a clear correlation between the χ metric (a controlled genetic distance method) with the dominance effects could be because we started from a single pool with clear LD patters as opposed to using genetic distance in natural populations where other subpopulation structure may affect the performance of genetic distance to predict dominance interactions. In summary, a big component of the total genetic value is the additive component, and the creation of controlled genetic distance through complementation should only be able to predict the dominance component of the equation but not the total performance as expected by previous studies in the 1980s and 1990s. Still, there is value in the genetic complementation as a way to create *de novo* heterotic pools quickly without phenotyping prior to start the GCA-based approaches.

The observation that at all levels of dominance, the complementation approach was shown to be almost as effective to increase the dominance values as selecting based on GCA is a promising result given the reduced complexity and resources (the only cost incurred is in genotyping a sample of the population in a nursery) involved in enabling the complementation approach for a couple of cycles before a GCA program is implemented. The observation that the complementation approach did not increase the additive value was not a surprise since no phenotyping is used in this method and there is no way to select for the positive allele. In a practical program the complementation approach would only be used to build dominance-based heterosis and drive the allele frequencies apart and then the GCA approach would need to come into play.

### Implementing genetic complementation in an active inbred program to start hybrid breeding

We recorded the additive, dominance and total genetic value of the parental lines and the advanced hybrids (in the case of the hybrid strategies) across 40 years of applying the different breeding strategies (Figure 6). The increase of performance of hybrids compared to lines when the simulated program moves from one pool to two pools but recycles parents based on BV (TWO_POOL_BV) is only due to the recovery of baseline heterosis rather than an increase in dominance or panmictic heterosis (based on the simulated level of dominance, meanDD=0.5) (Lamkey & Edwards, 1999; Labroo et al., 2021; Cowling at al. 2020). Increase in genetic gain in hybrids following a two pool BV method is based on the increase of additive gain.

Interestingly, the lower rate of increase in additive gain of RRS based on GCA compared to the TWO_POOL_BV programs is due to the additional time taken for the additional crossing and evaluation generation of testcrosses and the way the GCA estimate is constructed (Figure 6) (Hallauer et al., 2010). The lower rate of additive gain using GCA gets compensated by higher dominance gain if and only if the testers are updated (Figure 4). The decrease in additive value in lines and hybrids observed in the complementation approach (TWO_POOL_COMP) (Figure 6) is expected since both populations are trying to fix specific alleles without purging the negative alleles giving place to the average decrease in additive value due to inbreeding (Bernardo 1992, 2002). With incomplete dominance, the complementation approach the harnessing of heterosis comes at a “cost” to additive gain. In hybrids, such inbreeding is surpassed by the increase of dominance value when the mean dominance degree > 0.5, but as dominance variance gets depleted (in our simulations around year 15) inbreeding surpasses the effect of dominance value leading to stagnation of genetic gain in hybrids (Figure 6, Supplemental Figure 3).

### Similar performance with lower cost

As mentioned throughout this paper the major potential of the complementation methodology is the reduction of cost to produce high-performance hybrids in the long term through recurrent selection for dominance at a low cost when the species displays an intermediate or high level of mean dominance degree. The lowers cost of the complementation + RRS comes from the fact that the number of complementation cycles that the breeding program decides to run prior to start formal RRS only incurs in genotyping costs (∼$10 usd per sample) and a growing nursery (∼$5000 usd) where the material can be grown (without replication and experimental design) while the genotyping occurs and recombined once the complementation surrogate has been calculated. Assuming the same size than the simulated RRS program with 60 crosses and 50 F2s (3000 F2s) the complementation approach would give a cost of ∼ $35,000 usd per cycle. This cost per cycle is notable lower than a formal RRS cycle where a nursery (∼5000 usd), DH formation (∼$100K assuming ∼$2200 usd per cross producing ∼100 DHs per cross and 50 crosses), one per se evaluation of 6000 lines ($30K), a crossing block for testcross formation ($5000), and 3 seasons of testcross evaluation with 500, 200, 45 hybrids, (∼$30K per evaluation) without including other important costs.

A new complementation cycle can be immediately started since phenotyping for traits is not required, which allows to complete as many cycles in a year as the biological limit and speed breeding methods allow (Watson et al., 2018). In this paper we assumed a complementation cycle to last ∼1 year, but nothing impedes to use speed breeding to run 3 or more cycles per year and use 2-3 years of complementation prior to start formal RRS. Is important to highlight that the complementation cycles are just a preamble to start the formal RRS since the increase of additive gain and production of varieties requires the phenotyping and multi-environment testing that any breeding programs does, and the complementation approach can be seen as a boosting methodology to increase initially the dominance-based heterosis.

## Conclusion

The potential for hybrid breeding to fulfil the nutritional needs of a growing world population is based on the exploitation of heterosis, which is currently best explained by the theory of directional dominance. Hybrid breeding has conventionally been approached through general combining ability (GCA) using testcross-based or diallel-based approaches aiming to increase additive and dominance effects through recurrent selection, requiring substantial investment in phenotypic evaluation and a genomic selection variant to make gains faster. Here, we proposed that genetic marker data can be used to compute a surrogate of complementation between pools or subpopulations in order to accumulate dominance interactions through recurrent selection quickly and create *de novo* heterotic pools prior to start the GCA-based approaches. We found that the proposed genetic-complementation-based approach outperforms conventional approaches in terms of dominance gain per unit cost when dominance ranges from intermediate to high values (meanDD ≥ 0.5). The complementation approach can be thought as a coordinated genetic distance method or recurrent selection for dominance. The results in real datasets from dent by flint maize hybrids validate the complementation theory and surrogates of complementation. The complementation surrogate is an alternative for breeding programs attempting to transition from a non-hybrid to a hybrid system (in a diploid species) to increase and maximize the dominance-based heterosis per unit cost.

## Author Contributions

GCP, ML, CW, and DG conceived the study, GCP performed the simulations. Everyone contributed to writing the manuscript. All authors agree to be accountable for the content of the work.

## Acknowledgments

We thank the different breeding teams from the CGIAR that helped through their questions to conceive these ideas. We thank the reviewers for helping to improve the quality of this manuscript.

## Data availability statements

The datasets generated for this study can be found linked to the publications from Technow et al. (2014) and Kadam et al. (2016).

## Ethics approval

Not applicable.

## Consent to participate

Not applicable.

## Consent to publish

Not applicable.

**Supplementary Figure 1.**
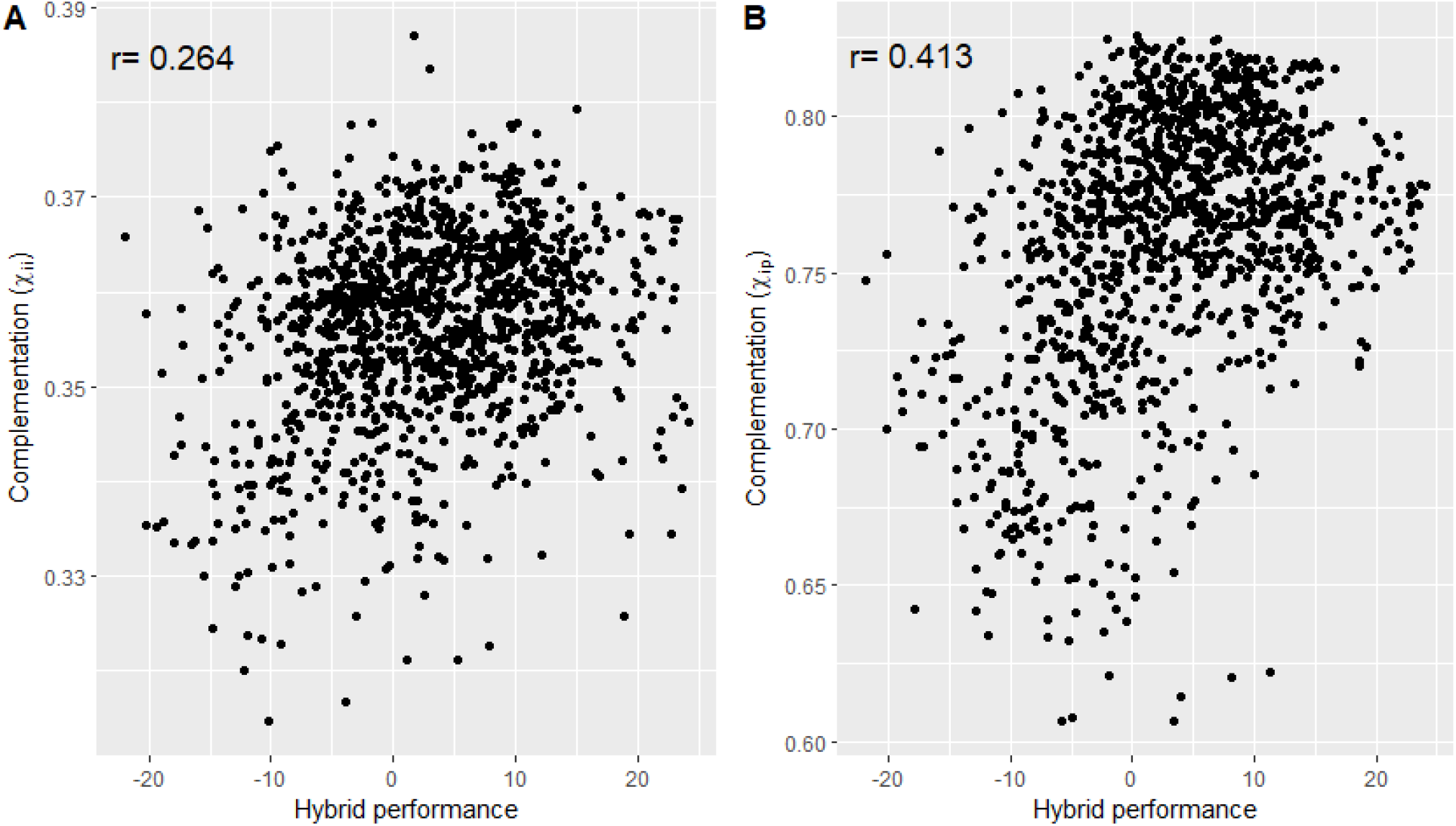
Genetic correlation between the complementation surrogates (individual-to-population and individual-to-individual complementation) and hybrid performance. The correlation legend is shown in the upper left corner of each plot. Data comes from Technow et al. (2014) and comprises a real dataset of hybrids between the dent and flint heterotic pools.

**Supplemental Figure 2.**
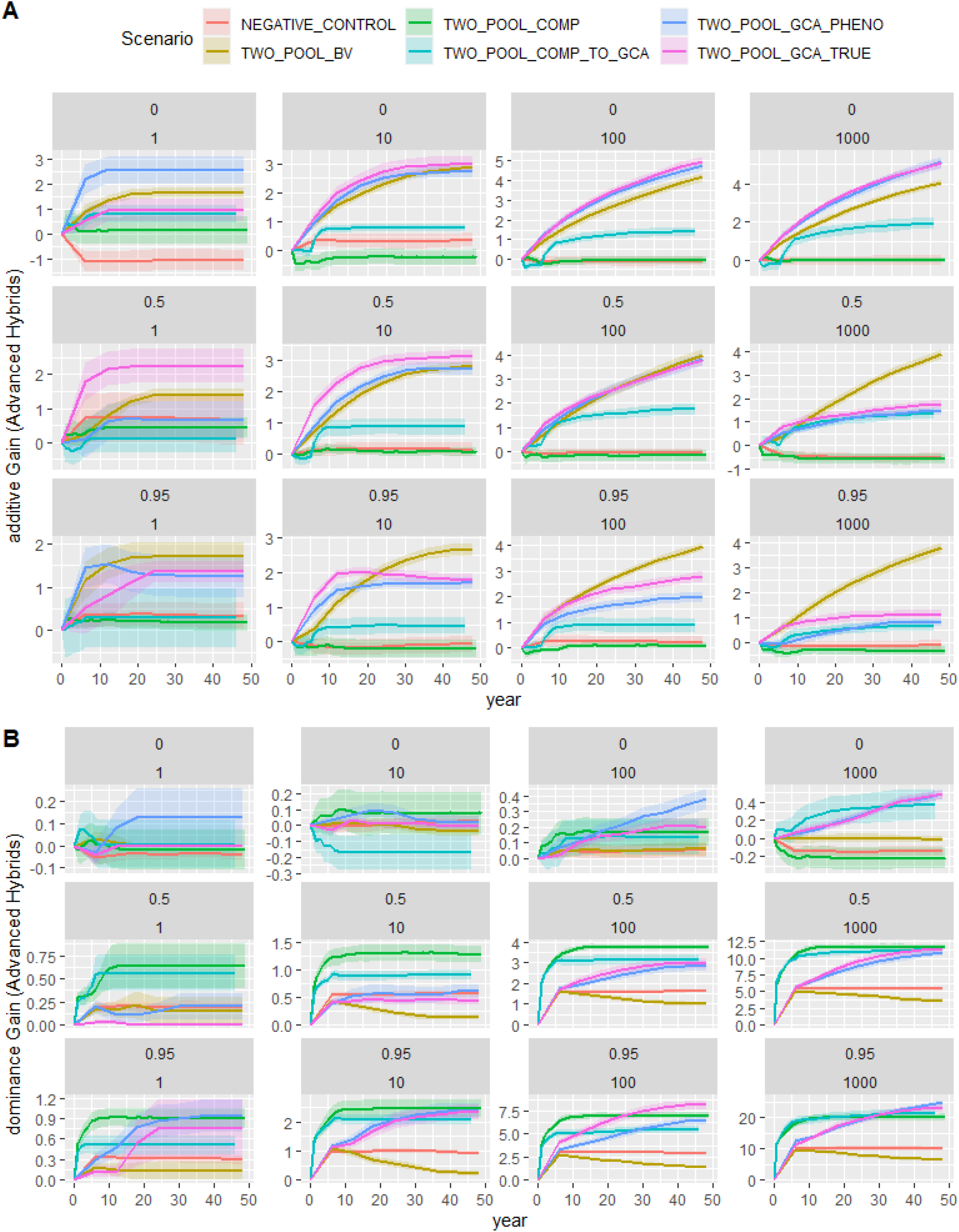
Additive and dominance for a trait in on-farm hybrids under different mean dominance degrees and number of QTLs for different simulated breeding strategies. The columns represent the comparison of strategies where the simulated trait has different number of QTLs behind (1, 10, 50, 100, 1000 QTLs). The rows represent the comparison of strategies where the simulated trait has different levels of mean dominance degree (0 implies a completely additive trait, 0.5 partial dominance, and 0.95 represents a trait with almost complete dominance). A value of varDD=0.2 was used across all scenarios. In the x axis, the number of selection cycles (20) are indicated whereas in the y axis the gain is shown. The different lines represent the different selection strategies: 1) reciprocal recurrent selection program that selects and recycles parents based on GCA (TWO_POOL_GCA_PHENO), 2) reciprocal recurrent selection program that selects and recycles parents based on genetic complementation (TWO_POOL_COMP), 3) reciprocal recurrent selection program that selects and recycles parents based on genetic complementation using true QTLs (TWO_POOL_COMP_TRUE; positive control 1), 4) reciprocal recurrent selection program that selects and recycles parents based on true GCA (TWO_POOL_GCA_TRUE; positive control 2), 5) recurrent selection program that selects and recycles parents based on breeding value (TWO_POOL_BV; negative control 1), 6) recurrent selection program that selects and recycles parents at random (ONE_POOL_RANDOM; negative control 2).

**Supplementary Figure 3.**
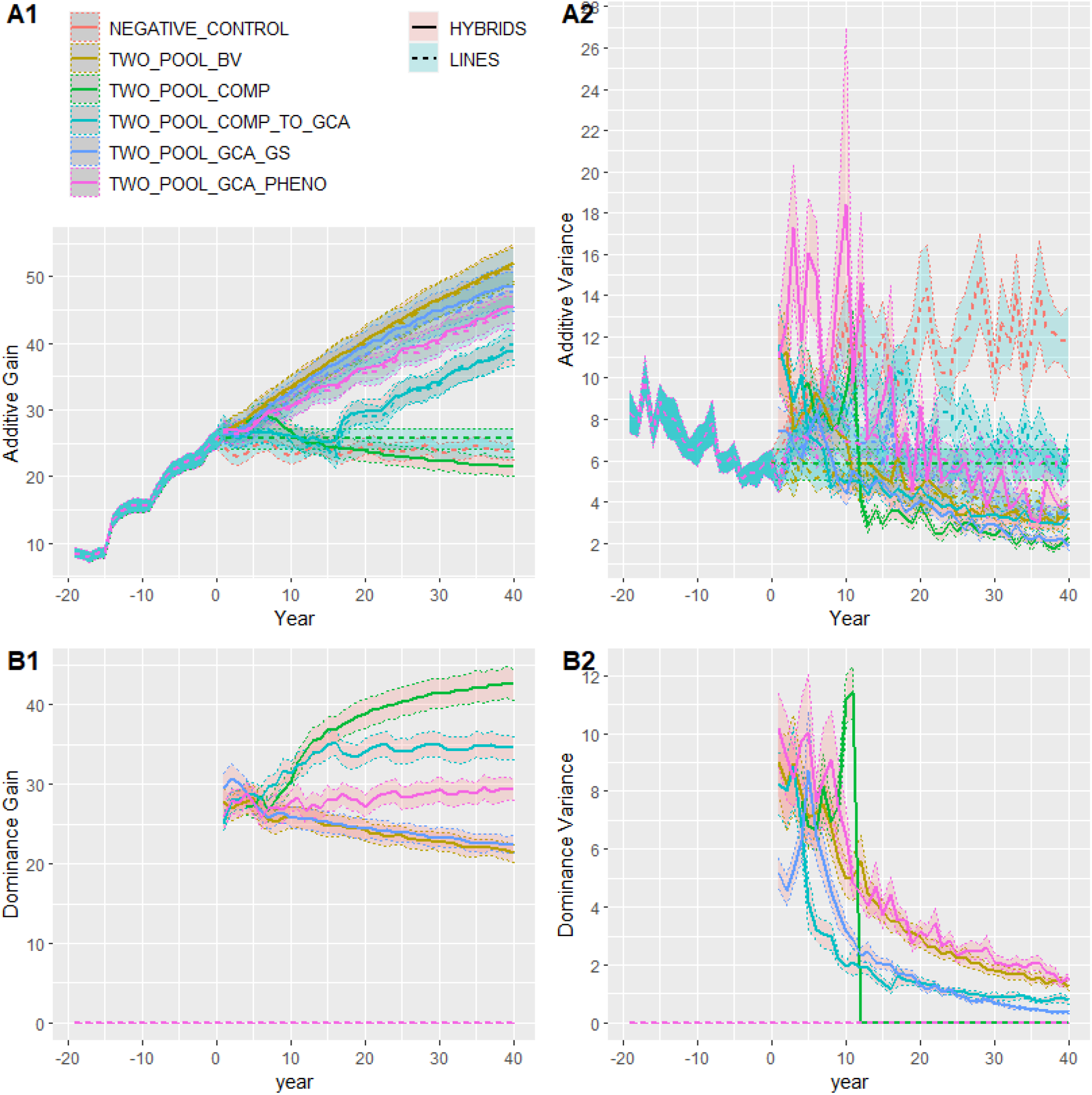
Genetic gain for a complex trait (1000 QTLs behind) with intermediate mean dominance degree (0.5) measured in on-farm hybrids (solid lines) and parental lines (dotted lines) for a simulated line program transitioning to a hybrid program with different breeding strategies. In the x axis, the number of years is indicated whereas in the y axis the genetic gain for (A) additive (B) dominance value are shown. The different lines represent the different selection strategies tested: 1) line program continuing the recurrent selection based on breeding value in two independent pools and making hybrids among the pools (TWO_POOL_BV; positive control), 2) line program transitioning to RRS to select and recycle parents based on GCA (TWO_POOL_GCA_PHENO), 3) line program transitioning to RRS to select and recycle parents based on predicted GCA using GS (TWO_POOL_GCA_GS), 4) line program transitioning to select purely on complementation (χ) (TWO_POOL_COMP), and 5) negative control selecting lines at random (NEGATIVE_CONTROL). Every year 4 out of the top 5 testers is updated. The value of varDD was 0.2 across all scenarios. Shadow lines represent standard errors for treatments across 20 Monte-Carlo replications.

## References

Allendorf, F. W. (1986). Genetic drift and the loss of alleles versus heterozygosity. Zoo biology, 5(2), 181–190.

Bernardo, R. (1992). Relationship between single-cross performance and molecular marker heterozygosity. Theoretical and Applied Genetics, 83(5), 628–634.

Bernardo, R. (2002). Breeding for quantitative traits in plants (Vol. 1, p. 369). Woodbury, MN: Stemma press.

Bingham, E. T. (1998). Role of chromosome blocks in heterosis and estimates of dominance and overdominance. Concepts and breeding of heterosis in crop plants, 25, 71–87.

Birchler, J. A., Yao, H., Chudalayandi, S., Vaiman, D., & Veitia, R. A. (2010). Heterosis. The Plant Cell, 22(7), 2105–2112.

Charcosset, A., Lefort-Buson, M., & Gallais, A. (1991). Relationship between heterosis and heterozygosity at marker loci: a theoretical computation. Theoretical and Applied Genetics, 81(5), 571–575.

Comstock, R. E., Robinson, H. F., & Harvey, P. H. (1949). Breeding procedure designed to make maximum use of both general and specific combining ability. Agronomy Journal.

Comstock, R. E., & Robinson, H. F. (1952). Estimation of average dominance of genes. Heterosis, 2, 494–516.

Crow, J. F. (1999). Dominance and overdominance. Genetics and exploitation of heterosis in crops, 49–58.

Cowling, W. A., Gaynor, R. C., Antolín, R., Gorjanc, G., Edwards, S. M., Powell, O., & Hickey, J. M. (2020). In silico simulation of future hybrid performance to evaluate heterotic pool formation in a self-pollinating crop. Scientific reports, 10(1), 1–8.

Davenport, C. B. (1908). Degeneration, albinism and inbreeding. Science, 28(718), 454–455.

de Boer, I. J., & Hoeschele, I. (1993). Genetic evaluation methods for populations with dominance and inbreeding. Theoretical and Applied Genetics, 86(2), 245–258.

Doebley, J. (2004). The genetics of maize evolution. Annu. Rev. Genet., 38, 37–59.

Dudley, J. W., Maroof, M. S., & Rufener, G. K. (1991). Molecular markers and grouping of parents in maize breeding programs. Crop Science, 31(3), 718–723.

Duvick, D. N., Smith, J. S. C., & Cooper, M. (2004). Long-term selection in a commercial hybrid maize breeding program. Plant breeding reviews, 24(2), 109–152.

East, E. M. (1936). Heterosis. Genetics, 21(4), 375.

Evenson, R. E., & Gollin, D. (2003). Assessing the impact of the Green Revolution, 1960 to 2000. science, 300(5620), 758–762.

Falconer, D.S., and T.F.C. Mackay. 1996. Introduction to quantitative genetics. 4th ed. Longman, Essex, England

Frei, O. M., Stuber, C. W., & Goodman, M. M. (1986). Use of Allozymes as Genetic Markers for Predicting Performance in Maize Single Cross Hybrids 1. Crop Science, 26(1), 37–42.

Gaynor, R. C., Gorjanc, G., & Hickey, J. M. (2021). AlphaSimR: an R package for breeding program simulations. G3, 11(2), jkaa017.

Godshalk, E. B., Lee, M., & Lamkey, K. R. (1990). Relationship of restriction fragment length polymorphisms to single-cross hybrid performance of maize. Theoretical and Applied Genetics, 80(2), 273–280.

Hallauer, A. R., Carena, M. J., & Miranda Filho, J. D. (2010). Quantitative genetics in maize breeding (Vol. 6). Springer Science & Business Media.

Hedden, P. (2003). The genes of the Green Revolution. TRENDS in Genetics, 19(1), 5–9.

Hill, W. G. (2010). Understanding and using quantitative genetic variation. Philosophical Transactions of the Royal Society B: Biological Sciences, 365(1537), 73–85.

Jiang, Y., Schmidt, R. H., Zhao, Y., & Reif, J. C. (2017). A quantitative genetic framework highlights the role of epistatic effects for grain-yield heterosis in bread wheat. Nature genetics, 49(12), 1741–1746.

Jinks, J. L., & Jones, R. M. (1958). Estimation of the components of heterosis. Genetics, 43(2), 223.

Joshi, P. K., Esko, T., Mattsson, H., Eklund, N., Gandin, I., Nutile, T., … & Kubo, M. (2015). Directional dominance on stature and cognition in diverse human populations. Nature, 523(7561), 459–462.

Kadam, D. C., Potts, S. M., Bohn, M. O., Lipka, A. E., & Lorenz, A. J. (2016). Genomic prediction of single crosses in the early stages of a maize hybrid breeding pipeline. G3: Genes, Genomes, Genetics, 6(11), 3443–3453.

Lande, R. (1976). Natural selection and random genetic drift in phenotypic evolution. Evolution, 314–334.

Lamkey, K. R., Hallauer, A. R., & Kahler, A. L. (1987). Allelic differences at enzyme loci and hybrid performance in maize. Journal of Heredity, 78(4), 231–234.

Lamkey, K. R., & Edwards, J. W. (1999). Quantitative genetics of heterosis.

Lee, M., Godshalk, E. B., Lamkey, K. R., & Woodman, W. W. (1989). Association of restriction fragment length polymorphisms among maize inbreds with agronomic performance of their crosses. Crop Science, 29(4), 1067.

Lippman, Z. B., & Zamir, D. (2007). Heterosis: revisiting the magic. Trends in genetics, 23(2), 60–66.

Lynch, M., & Walsh, B. (1998). Genetics and analysis of quantitative traits.

Lynch, M., Ackerman, M. S., Gout, J. F., Long, H., Sung, W., Thomas, W. K., & Foster, P. L. (2016). Genetic drift, selection and the evolution of the mutation rate. Nature Reviews Genetics, 17(11), 704–714.

Melchinger, A. E., Lee, M., Lamkey, K. R., & Woodman, W. L. (1990). Genetic diversity for restriction fragment length polymorphisms: relation to estimated genetic effects in maize inbreds. Crop science, 30(5), 1033.

Mrode, R. A. (2014). Linear models for the prediction of animal breeding values. Cabi.

Nishio, M., & Satoh, M. (2014). Including dominance effects in the genomic BLUP method for genomic evaluation. PloS one, 9(1), e85792.

Rembe, M., Zhao, Y., Jiang, Y., & Reif, J. C. (2019). Reciprocal recurrent genomic selection: an attractive tool to leverage hybrid wheat breeding. Theoretical and Applied Genetics, 132(3), 687–698.

Schnell, F. W. (1965). Die Covarianz zwischen Verwandten in einer gen-orthogonalen Population. I. Allgemeine Theorie. Biometrische Zeitschrift, 7(1), 1–49.

Su, G., Christensen, O. F., Ostersen, T., Henryon, M., & Lund, M. S. (2012). Estimating additive and non-additive genetic variances and predicting genetic merits using genome-wide dense single nucleotide polymorphism markers.

Varona, L., Legarra, A., Herring, W., & Vitezica, Z. G. (2018). Genomic selection models for directional dominance: an example for litter size in pigs. Genetics Selection Evolution, 50(1), 1–13.

Watson, A., Ghosh, S., Williams, M. J., Cuddy, W. S., Simmonds, J., Rey, M. D., … & Hickey, L. T. (2018). Speed breeding is a powerful tool to accelerate crop research and breeding. Nature plants, 4(1), 23–29.

Werner, C. R., Gaynor, R. C., Sargent, D. J., Lillo, A., Gorjanc, G., & Hickey, J. M. (2020). Genomic selection strategies for clonally propagated crops. bioRxiv.

Zhang, Q., Zhou, Z. Q., Yang, G. P., Xu, C. G., Liu, K. D., & Maroof, M. S. (1996). Molecular marker heterozygosity and hybrid performance in indica and japonica rice. Theoretical and applied genetics, 93(8), 1218–1224.

Zhao, Y., Li, Z., Liu, G., Jiang, Y., Maurer, H. P., Würschum, T., … & Reif, J. C. (2015). Genome-based establishment of a high-yielding heterotic pattern for hybrid wheat breeding. Proceedings of the National Academy of Sciences, 112(51), 15624–15629.

