## Supplementary figures and images for "Reciprocal recurrent selection based on genetic complementation: An efficient way to build heterosis in diploids due to directional dominance"

### Supplemental Figure 1

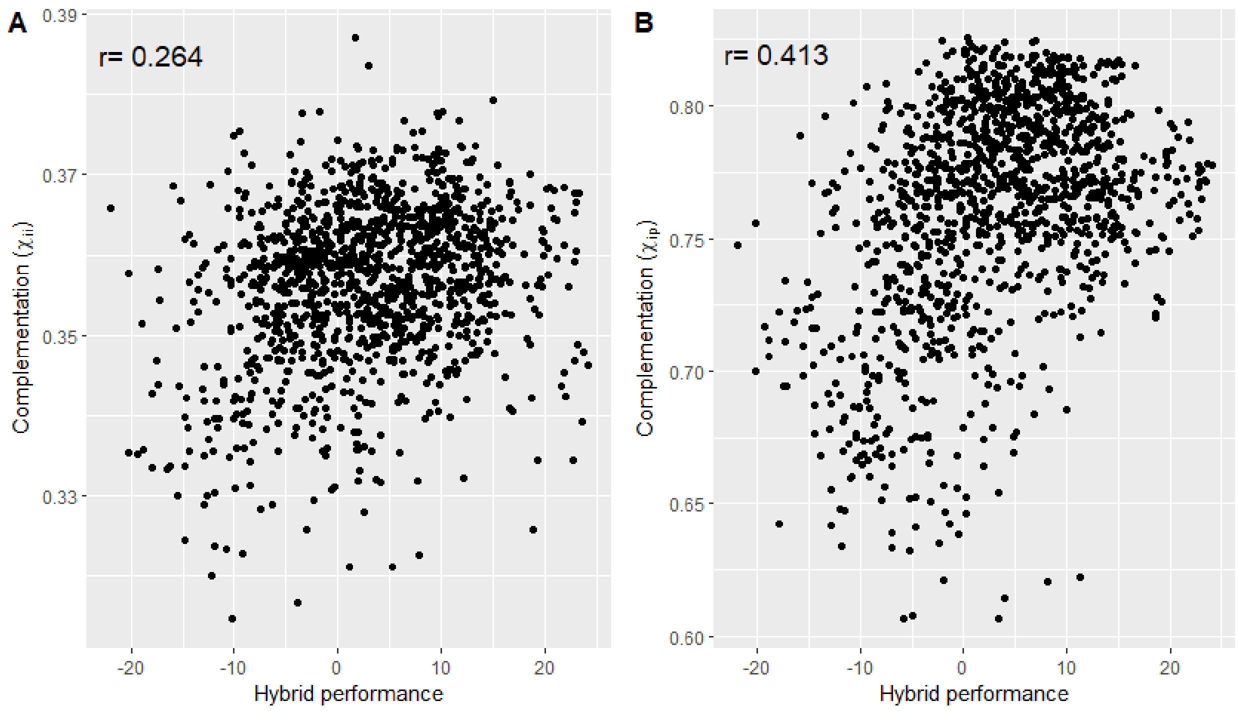

### Supplemental Figure 2

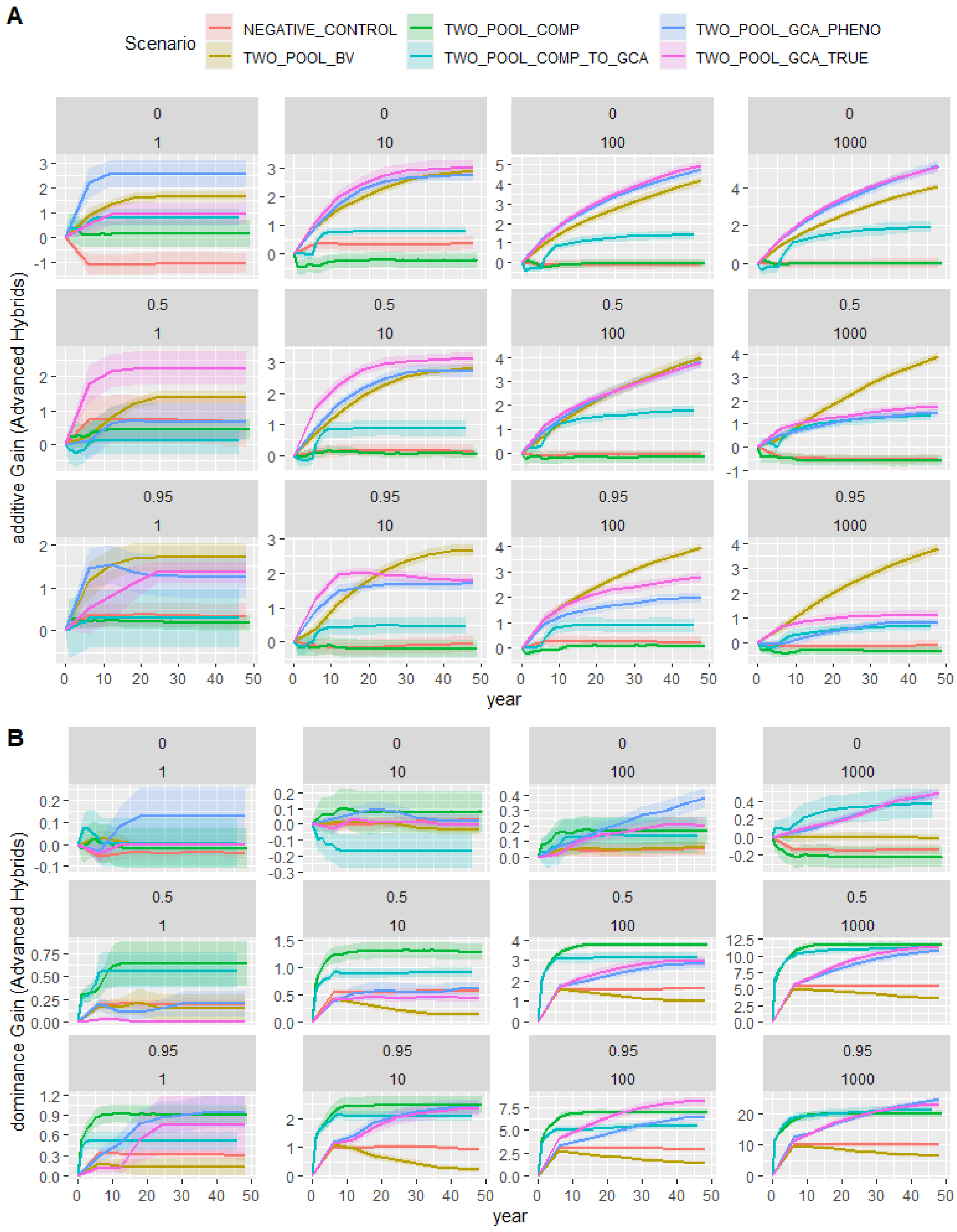

### Supplemental Figure 3

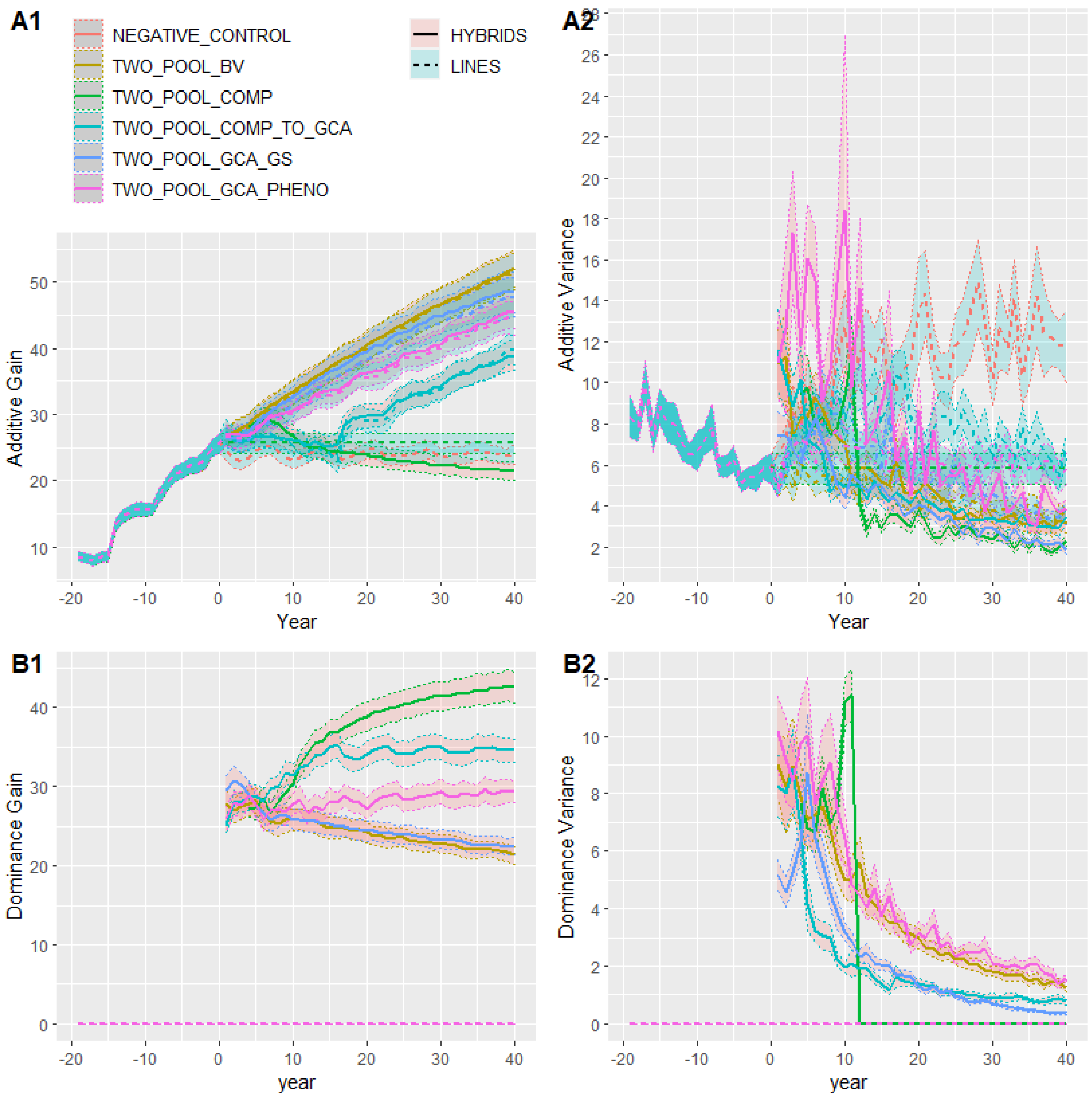
